# Genome-wide identification of the LexA-mediated DNA damage response in *Streptomyces venezuelae*

**DOI:** 10.1101/2022.03.28.486056

**Authors:** Kathryn J. Stratton, Matthew J. Bush, Govind Chandra, Clare E. M. Stevenson, Kim C. Findlay, Susan Schlimpert

## Abstract

DNA damage triggers a widely conserved stress response in bacteria called the SOS response that involves two key regulators, the activator RecA and the transcriptional repressor LexA. Despite the wide conservation of the SOS response, the number of genes controlled by LexA varies considerably between different organisms. The filamentous soil-dwelling bacteria of the genus *Streptomyces* contain LexA and RecA homologs but their roles in *Streptomyces* have not been systematically studied. Here, we demonstrate that RecA and LexA are required for the survival of *Streptomyces venezuelae* during DNA damaging conditions and for normal development during unperturbed growth. Monitoring the activity of a fluorescent *recA* promoter fusion and LexA protein levels revealed that the activation of the SOS response is delayed in *S. venezuelae*. By combining global transcriptional profiling and ChIP-seq analysis, we determined the LexA regulon and defined the core set of DNA damage repair genes that are expressed in response to treatment with the DNA alkylating agent mitomycin C. Our results show that DNA damage-induced degradation of LexA results in the differential regulation of LexA target genes. Using Surface Plasmon Resonance, we further confirm the LexA DNA binding motif (SOS box) and demonstrate that LexA displays tight but distinct binding affinities to its target promoters, indicating a graded response to DNA damage.

**IMPORTANCE:** The transcriptional regulator LexA functions as a repressor of the bacterial SOS response, which is induced during DNA damaging conditions. This results in the expression of genes important for survival and adaptation. Here, we report the regulatory network controlled by LexA in the filamentous antibiotic producing *Streptomyces* bacteria and establish the existence of the SOS response in *Streptomyces*. Collectively, our work reveals significant insights into the DNA damage response in *Streptomyces* that will promote further studies to understand how these important bacteria adapt to their environment.

## INTRODUCTION

Exposure to genotoxic agents can cause DNA damage with potentially lethal consequences and therefore the ability to restore genome integrity is an essential task for all living organisms. In bacteria, DNA damage triggers a widely conserved cellular stress response called the SOS response (1). The SOS response is regulated by two key proteins, the activator RecA (a multifunctional recombinase) and the transcriptional repressor LexA. During unperturbed growth, LexA binds as a dimer to a 16-bp palindromic sequence motif, the SOS box, located in the promoter region of its target genes, thereby preventing their expression. Elevated levels of DNA damage can lead to the accumulation of single-stranded (ss) DNA, which is sensed and bound by RecA, resulting in the formation of a nucleoprotein filament. This activated form of RecA functions as a co-protease and promotes the self-cleavage of LexA. Consequently, LexA repressor levels decrease in the cell, leading to the induction of the SOS response regulon. The gene products of the SOS regulon are involved in a range of cellular functions critical for survival, adaptation, and DNA repair (2). Specifically, SOS induction triggers the expression of genes involved in nucleotide excision and recombination repair (*uvrABD, recN, ruvCAB*), the removal of alkylation damage (*ada, alkAB, aidB*) and translesion synthesis (*polB, dinP, umuDC, dnaE2*) (3). In addition, the LexA regulon often activates species-specific cell cycle checkpoints that stall cellular development to allow sufficient time for DNA repair to occur (4). LexA also negatively autoregulates its own transcription and *recA* expression (5), thereby ensuring that the SOS response is rapidly turned off once genome integrity has been restored.

The two principal regulators of the SOS response, RecA and LexA, are evolutionary well conserved and present in almost all free-living bacteria, including the ubiquitous Gram-positive Actinomycetota (previously Actinobacteria) (3). Prominent members of this phylum are the streptomycetes, which are renowned for their extraordinary potential to produce specialised metabolites of economic and medical importance (6). *Streptomyces* are characterized by a sophisticated developmental life cycle that starts with the germination of spores. Emerging germ tubes grow by apical tip extension and hyphal branching to establish a dense vegetative mycelial network that scavenges for nutrients. In response to nutrient starvation, hyphae escape surface tension and grow upwards to form so-called aerial hyphae. These hyphae subsequently undergo reproductive cell division, resulting in the production of chains of equally-sized unigenomic spores that are eventually released into the environment (7).

*Streptomyces* are commonly found in sediments, a heterogeneous and competitive habitat where they are constantly exposed to a range of environmental stressors, including genotoxic agents. Earlier studies showed that a *Streptomyces recA* null mutant is more sensitive to DNA damaging agents and that DNA damaging antibiotics can inhibit sporulation, suggesting the existence of an SOS-like stress response (8, 9). However, the underlying regulatory network that underpins such a stress response is not understood. Furthermore, work over the last two decades has revealed that the SOS motif recognized by LexA and the number of genes directly controlled by LexA vary between bacterial genera and cannot simply be predicted bioinformatically (2, 3).

In this work we confirm the existence of the SOS response in *Streptomyces venezuelae*, one of the key model organisms for this genus (7, 10), and demonstrate that RecA and LexA are important for survival during DNA damaging growth conditions. By combining global transcriptomic profiling and ChIP-seq analysis, we identify the LexA regulon and the core set of SOS genes involved in DNA repair. Using Surface Plasmon Resonance, we determine the LexA binding kinetics to selected target promoters and confirm the specific interaction of LexA with the identified SOS box. Our *in vivo* and *in vitro* work further reveals that in *S. venezuelae* LexA displays a differential DNA binding affinity that implies a fine-tuned activation of the SOS regulon in response to DNA damage.

## RESULTS

### Deletion of *recA* and *lexA* affects sporulation and sensitizes *S. venezuelae* to DNA damage

*The S. venezuelae* genome contains a single *recA* gene (*vnz_26845*) and a single *lexA* gene (*vnz_27115*). To characterize the SOS response in *S. venezuelae*, we first constructed a Δ*recA* mutant by replacing the *recA* coding region with an apramycin resistance cassette. Deletion of *recA* resulted in a growth defect and the formation of two colony types: minute nonviable colonies and slow-growing colonies with delayed and irregular sporulation septation. A similar Δ*recA* phenotype was previously described for *S. coelicolor* (8). The developmental defects of the Δ*recA* mutant could be fully complemented by expressing *recA in trans* from its native promoter (Figure 1A and Supplementary Figure 1A). To exclude the possibility that a compensating mutation allowed for more robust growth in the slow-growing Δ*recA* mutant, we sequenced the genomic DNA isolated from two representative colonies and the parental strain. We did not detect a suppressor mutation in the Δ*recA* mutant, suggesting that both colony phenotypes are a direct consequence of the *recA* deletion.

**Figure 1:**
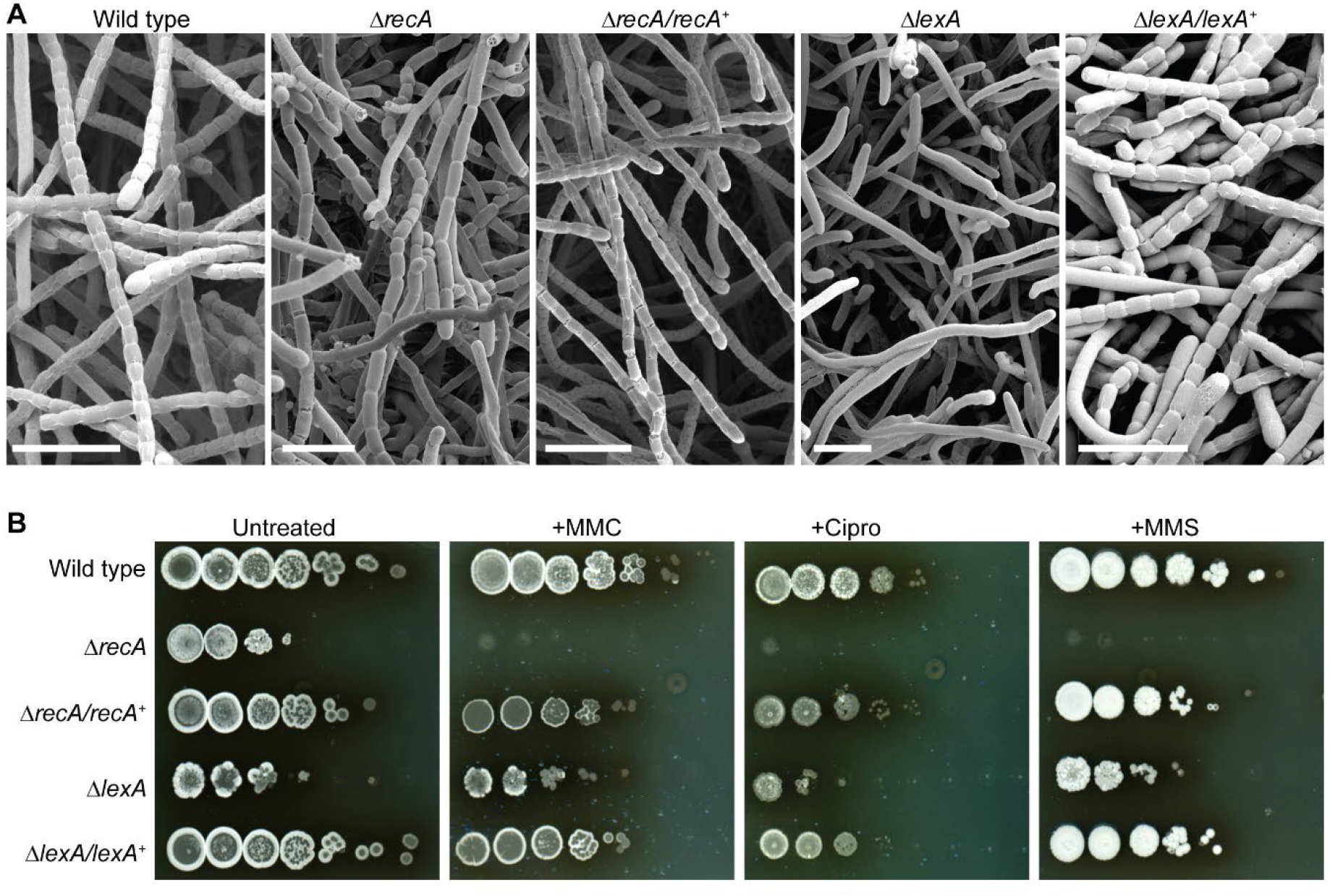
Deletion of *recA* and *lexA* impair cellular development and increase sensitivity to DNA damaging agents in *S. venezuelae*. **A)** Representative cryo-scanning electron micrographs showing aerial hyphae from the wild type, the Δ*recA* mutant (KS3), the Δ*lexA* mutant (KS44) and complemented mutant strains Δ*recA/recA*^+^ (KS14) and Δ*lexA/lexA*^+^ (KS57). Scale bars: 5 μm. **B)** Viability of the same strains as above when grown on solid MYM in the absence and presence of 0.25 μg ml^-1^ mitomycin C (MMC), 3 μg ml^-1^ ciprofloxacin (Cipro) and 3 mM methyl methanesulfonate (MMS). Equal concentrations of spores were used, and a 10-fold serial dilution series of the spore suspension was spotted onto MYM agar and incubated for 3-4 days. Shown are representative images from biological triplicate experiments.

Next, we generated a *S. venezuelae lexA* null mutant. Using a three-step procedure, a second copy of *lexA* was first integrated at the phage BT1 attachment site of wild-type *S. venezuelae*, followed by the deletion of the native *lexA* gene and the insertion of an apramycin resistance cassette. The apramycin-marked Δ*lexA* allele was then moved back into wild-type *S. venezuelae*, which lacked a second copy of *lexA*, using generalized SV1 phage transduction, generating the Δ*lexA* mutant strain. Notably, deletion of *lexA* resulted in a severe growth defect and a “white” phenotype (Supplementary Figure 1A), indicating that sporulation was impaired in this genetic background. Scanning electron micrographs of colony surfaces confirmed that the Δ*lexA* mutant failed to deposit regularly spaced sporulation septa, resulting in undifferentiated aerial hyphae and virtually no spores (Figure 1A). Normal sporulation and pigmentation were restored to the Δ*lexA* mutant when a single copy of wild-type *lexA* under the control of its native promoter was introduced *in trans* (Figure 1A and Supplementary Figure 1A).

We then determined the viability of the wild type, the Δ*recA* mutant, the Δ*lexA* mutant and the corresponding complemented mutant strains in the presence of DNA damaging agents, including mitomycin C (MMC), ciprofloxacin (Cipro) and methyl methanesulfonate (MMS). MMC and MMS are DNA alkylating agents that cause DNA double-strand breaks while ciprofloxacin targets DNA gyrase and inhibits DNA replication, leading to the accumulation of single-stranded DNA. For this we either performed multiwell-based liquid growth assays or spotted serial dilutions of the individual strains onto solid medium supplemented with MMC, ciprofloxacin or MMS. Based on our initial liquid growth assays we found that treatment of wild-type *S. venezuelae* with 0.25 µg ml^-1^ MMC, 3 µg ml^-1^ ciprofloxacin, and 3-5 mM MMS, strongly inhibited growth (Figure 1B and Supplementary Figure 1B). As expected, the absence of either *recA* or *lexA* dramatically reduced the viability of *S. venezuelae* when grown in the presence of MMC, ciprofloxacin or MMS and thus, as in other bacteria, RecA and LexA are essential for the survival of genotoxic stress in *S. venezuelae*.

### Induction of the SOS response requires prolonged exposure to DNA damaging agents

The *Streptomyces* life cycle includes a multicellular growth phase during which growing multinucleoid filaments are segmented by cross-walls. To gain an initial insight into the spatial and temporal activation of the SOS response within the mycelium, we constructed a strain in which expression of an ectopically inserted copy of *mcherry* was driven by the *recA* promoter and monitored the increase of mCherry fluorescence in response to MMC-induced DNA damage. We found that exposure of vegetative growing *S. venezuelae* cells to 0.25 µg ml^-1^ MMC for 1h or 2h did not lead to marked activation of the P_*recA*_*-mcherry* reporter fusion. However, when *S. venezuelae* was grown in the presence of MMC for 14h, a clear increase in cytoplasmic mCherry fluorescence within the entire mycelium was detectable (Figure 2A). Notably, we obtained similar results when treating *S. venezuelae* with 3 μg ml^-1^ ciprofloxacin or 3 mM methyl methansulfonate for 1h or 14h, respectively, indicating that the delayed induction of *recA* expression was not specific to mitomycin C (Supplementary Figure 2A). We also repeated the live cell imaging experiments using a 10-fold higher MMC concentrations (2.5 µg ml^-1^) to test if this would trigger a more rapid induction of the SOS response but did not detect increased expression of the *recA-mcherry* promoter fusion (Figure 2B).

**Figure 2:**
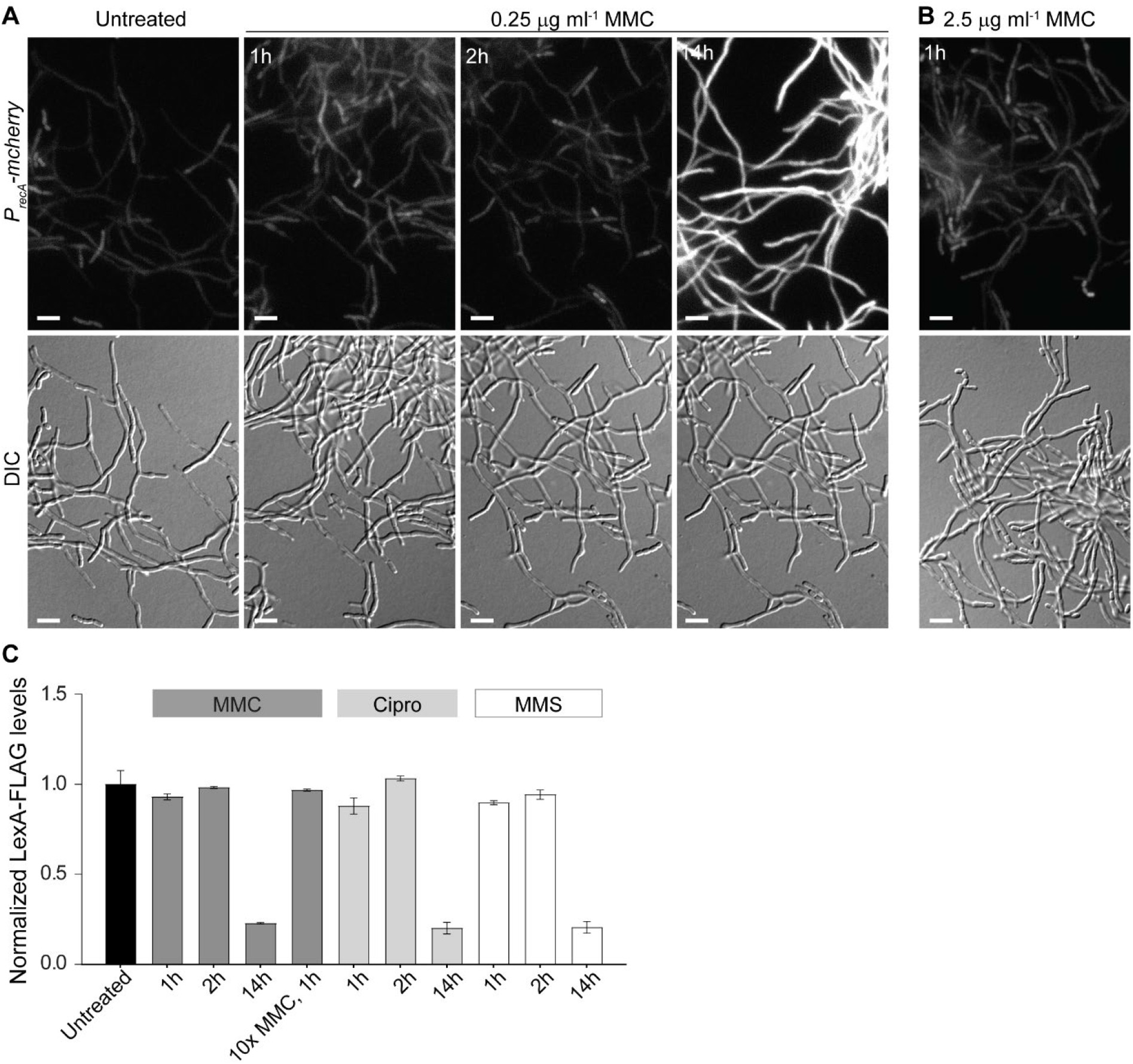
Induction of the SOS response requires prolonged exposure to DNA damaging agents. **A-B)** Fluorescence micrographs and corresponding differential interference contrast (DIC) images showing the induction of the *recA-mcherry* promoter fusion (*P*_*recA*_*-mcherry*) in wild-type *S. venezuelae* hyphae (KS80) grown in the absence and presence of **(A)** 0.25 μg ml^-1^ mitomycin C (MMC) for 1, 2 and 14h or **(B)** with 2.5 μg ml^-1^ mitomycin C for 1h. Shown are representative images from biological triplicate experiments. Scale bars: 5 μm. **C)** Automated western blot analysis of LexA-FLAG protein levels in *S. venezuelae* (Δ*lexA/lexA-FLAG*^*+*^, KS74) grown without and with 0.25 μg ml^-1^ and 2.5 μg ml^-1^ MMC, 3 μg ml^-1^ ciprofloxacin (Cipro) and 5 mM methyl methanesulfonate (MMS) for 1h, 2h and 14h. LexA-FLAG abundance was determined using a monoclonal anti-FLAG antibody. Protein levels were normalized to the untreated control samples. Analysis was performed in biological duplicate. Representative virtual Western blots are shown in Supplementary Figure 4A.

To corroborate these findings, we engineered a functional LexA-FLAG fusion that complemented the growth and sporulation defect in hyphae lacking *lexA* (Supplementary figure 2B). Vegetatively growing cultures expressing a single copy of *lexA* fused to the *3xFLAG* tag were challenged with 0.25 µg ml^-1^ MMC for 1h, 2h or 14h and subsequently, LexA-FLAG abundance in the corresponding cell lysates was analysed by automated Western blotting. In line with fluorescent *recA* reporter fusion results (Figure 2A), the LexA-FLAG protein level remained largely stable following treatment with MMC for 1 and 2 hours. However, LexA-FLAG was almost completely absent in cell lysates of samples grown in the presence of MMC for 14h (Figure 2C). To test the possibility that the FLAG tag was responsible for the slow degradation of LexA-FLAG fusion, we used a polyclonal LexA antibody to examine the decrease in native LexA levels in wild-type *S. venezuelae* following the same MMC treatment. We observed the same slow decrease in native LexA levels as we saw in the strain producing the LexA-FLAG fusion, showing that the FLAG tag did not affect the rate of degradation of LexA (Supplementary Figure 2C). Together, these results indicate that the activation of the two key components involved in launching the SOS response requires a chronic exposure to genotoxic stress in *Streptomyces*.

### Defining the LexA DNA damage response regulon

To identify genes directly under the control of LexA, we performed chromatin immunoprecipitation sequencing (ChIP-seq), using anti-FLAG antibody on the Δ*lexA* strain that expressed a functional LexA-FLAG fusion (Supplementary Figure 2C). Wild-type *S. venezuelae* was used as a negative control to eliminate any false signals that might arise from cross-reaction of the anti-FLAG antibody with other DNA binding proteins. ChIP-seq revealed numerous LexA binding sites distributed across the *S. venezuelae* genome in the absence of MMC (Figure 3A). No significant enrichment was observed in the wild-type control, suggesting that there was no cross-reaction of the anti-FLAG antibody leading to non-specific enrichment (Figure 3B). In total, 498 ChIP-seq peaks were identified, corresponding to LexA binding sites that were enriched more than 2-fold relative to the wild-type control. Of these, we selected all the binding sites that were located within a potential regulatory region of - 200 to 100 bp relative to the start codon of the first gene in a transcriptional unit (Supplementary Table 1). Through this route, we determined 270 putative LexA binding sites.

**Figure 3:**
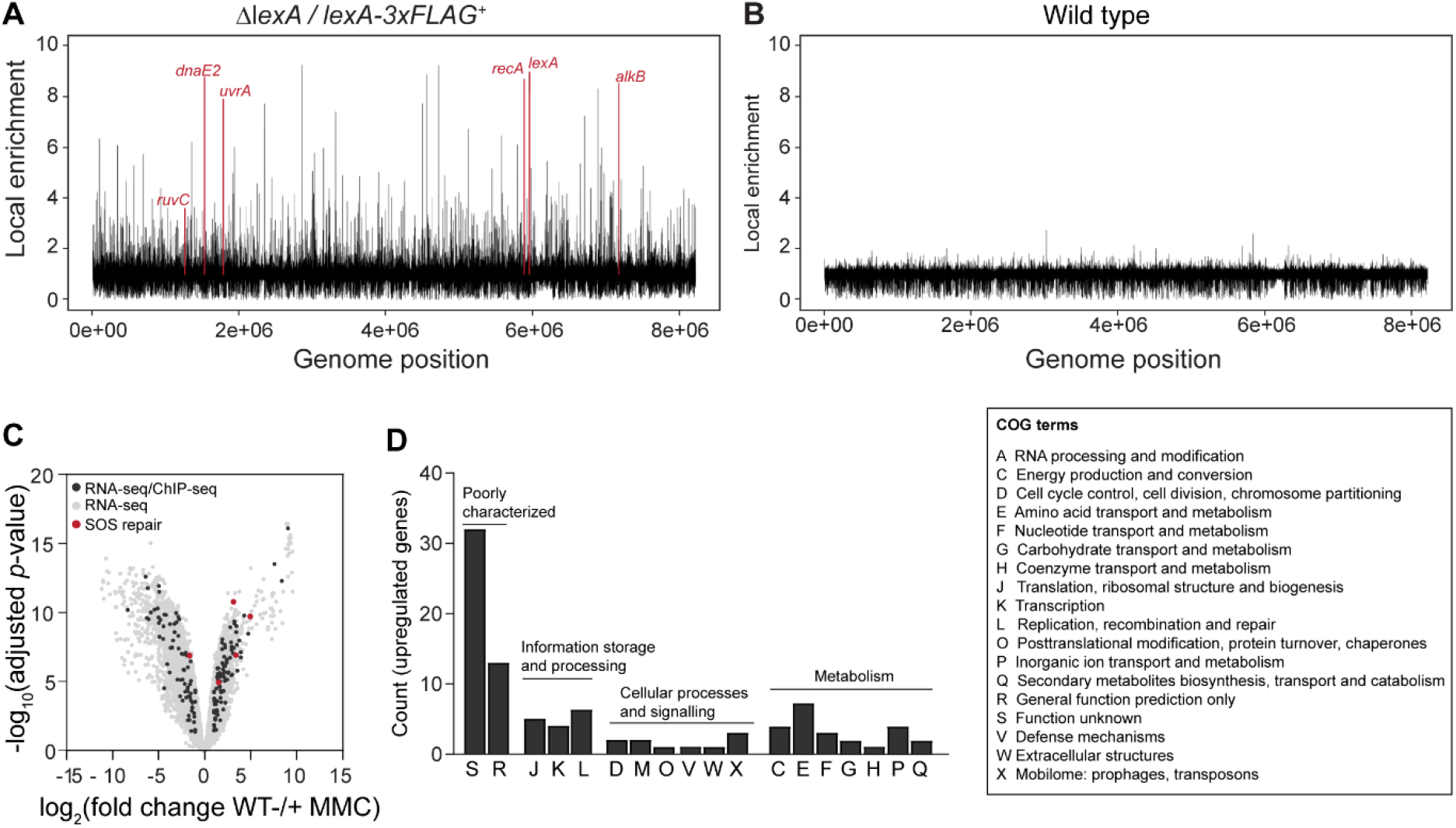
Genome-wide identification of the LexA regulon. **(A-B)** Genome-wide ChIP peak distribution of the LexA regulon in the **(A)** Δ*lexA/lexA-FLAG*^*+*^ strain (KS74) and **(B)** the wild type. Strains were grown for 14h in MYM. Peaks of selected LexA regulon genes involved in DNA repair are labelled and coloured in red lines. ChIP-seq results were obtained from biological replicate experiments. **(C)** Volcano plot comparing RNA-seq profiles of the wild type grown with or without 0.25 μg ml^-1^ MMC for 14h. Differentially expressed genes (grey dots) are defined by the adjusted *p*-value (*p*<0.05) and a fold change cut-off (|log2-fold change|>1). LexA target genes identified by ChIP-seq are labelled in black (n=175). Red-labelled dots indicate core DNA repair genes in the SOS regulon (Table 1). RNA-seq data was obtained from biological triplicate experiments. **(D)** Functional classification of genes that were upregulated following the degradation of LexA in response to MMC-induced stress (n=97, log2-fold change>1, adjusted *p*-value<0.05) were assigned Cluster of Orthologous Groups (COG) identifiers and their distribution was plotted. A legend for the corresponding COG annotations is presented to the right of the graph. Source data used to generate panel A-D can be found in Supplementary Table 1,3 and 4.

Furthermore, we detected 176 intragenic LexA binding sites that were in proximity of previously mapped transcription start sites (11), indicating that LexA might also control the activity of intragenic promoters. We note that the transcription start site mapping was performed in the absence of DNA damage, a condition during which LexA remains bound to its target operators and largely prevents transcription of the associated genes. Closer inspection of the putative intragenic promoters associated with LexA enrichment sites revealed, that six out of the 176 putative intragenic promoters were located within genes with a proposed function in DNA replication and repair (vnz_05400, vnz_06010, 08430, vnz_25520, vnz_29285) and within one gene that shares similarity with the cell division protein ZapE (*vnz_29020*) (12). In addition, two noticeable intragenic LexA binding sites for which no transcription start site data is available were detected within the DNA helicase *uvrD* (*vnz_23895*) and the error-prone DNA polymerase *dnaE2* (*vnz_06640*) (Supplementary Table 2). However, with the exception of dnaE2, none of the genes associated with an intragenic LexA binding sites were differentially regulated in response to DNA damage (Supplementary Table 3).

In order to identify LexA target genes that showed a clear transcriptional response to DNA damage, we performed RNA sequencing on wild-type *S. venezuelae* treated with 0.25 µg ml^-1^ MMC, combined with an untreated wild-type control. Because we only observed significant induction of *recA* expression and LexA degradation after prolonged exposure to MMC, we conducted RNA-seq in three independent biological samples obtained from cultures that were grown in the presence and absence of MMC for 14h. Due to the severely reduced viability of the Δ*recA* mutant when grown in medium containing MMC (Figure 1B), we were unable to include this strain in the RNA-seq analysis. In this way, we were able to identify 175 LexA binding sites that were associated with differentially expressed transcriptional units upon treatment with MMC (adjusted *p-*value *p*<0.05, |log2-fold change|>1) (Figure 3C and Supplementary Table 3). Of these, 97 transcriptional units were upregulated, including one putative LexA-target of unknown function located on the *S. venezuelae* plasmid pSVIJ1 and 78 transcriptional units were downregulated following MMC treatment (10). Although LexA generally functions as transcriptional repressor (13–15), the MMC-induced downregulation of genes associated with a LexA binding site is indicative of a direct or indirect positive regulation of these genes by LexA. We note that our global transcriptional profiling analysis further showed that several genes required for sporulation were significantly downregulated following the treatment with MMC, including the classical *Streptomyces* developmental regulatory genes *whiI, whiH* and *whiB*, genes involved in cell division *(ssgB)* and chromosome segregation *(smeA-sffA)* and several genes within the *whiE* locus, which encodes the biosynthesis machinery for the production of the polyketide pigment associated with mature spores (16–19). These findings are in agreement with other recent studies, reporting that subinhibitory concentrations of MMC inhibit sporulation in *S. venezuelae* (9, 20).

Functional categorization of the 97 LexA target genes derepressed in response to MMC stress did not suggest an enrichment of a specific gene category and revealed that the function of most genes associated with a LexA binding site is unknown and (Figure 3D and Table 1). Among the LexA targets that showed the highest upregulation were several genes belonging to a predicted prophage cluster (*vnz_21525, vnz_21545, vnz_21550*), genes functioning in protein translation and degradation (*vnz_05845* and *vnz_00275*), metabolism (*vnz_22770*), molecule transport (*vnz_15995 and vnz_26265*). Importantly, our integrated RNA-seq and ChIP-seq approach led to the identification of a core set of LexA targets (3), including the *recA-recX operon, lexA* and several conserved DNA repair genes encoding the translesion DNA polymerases DnaE2 and DinP, RuvC and RuvA required for recombination and repair, and the alkylation repair protein AlkB. In addition, our data suggest that *uvrA* (nucleotide excision repair) may also be part of the LexA-controlled DNA damage response as we observed a clear LexA ChIP-seq peak and a change in *uvrA* expression although this was just beneath our applied cut-off in the RNA-seq analysis (|log2-fold change|>1).

**Table 1:**
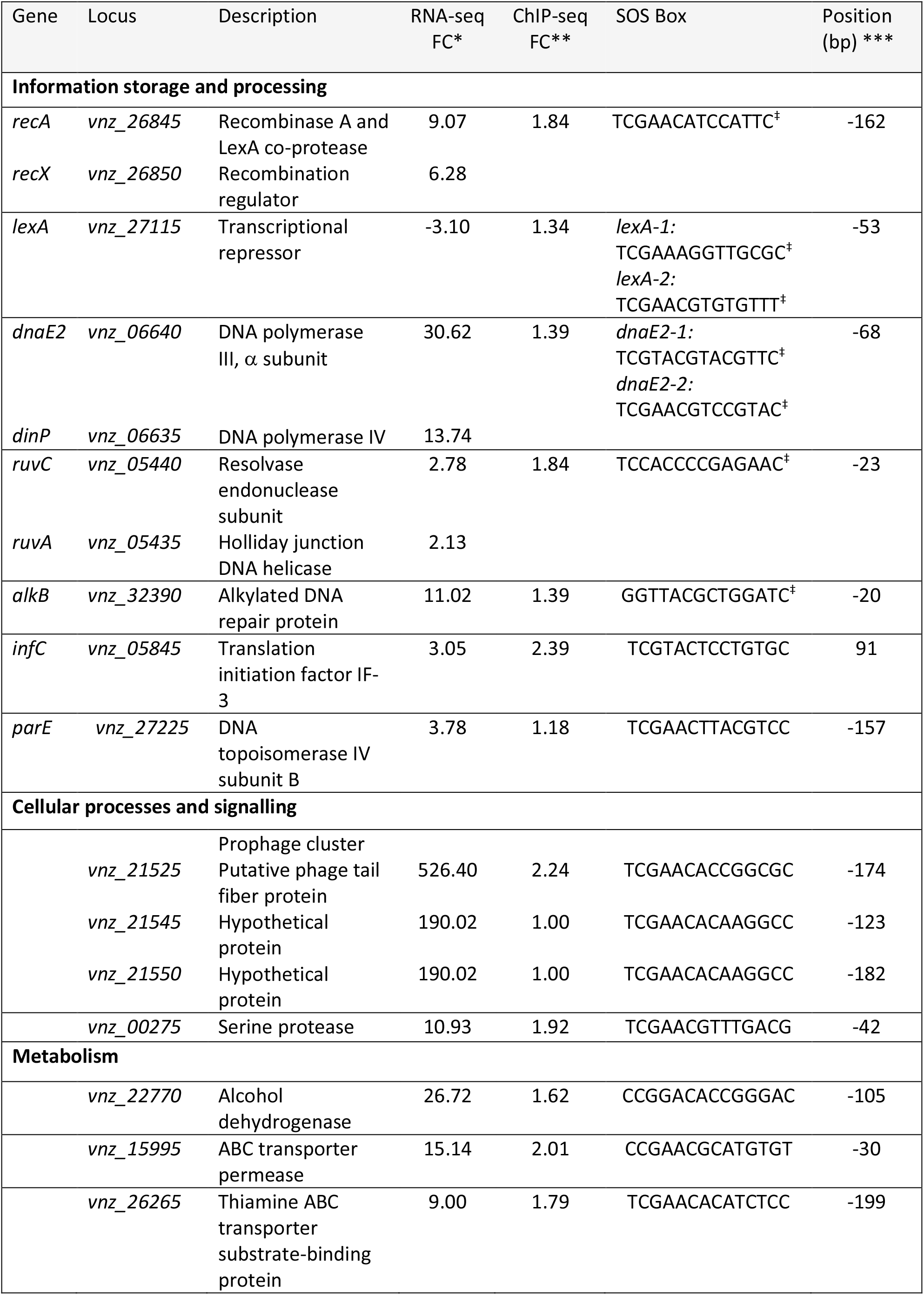
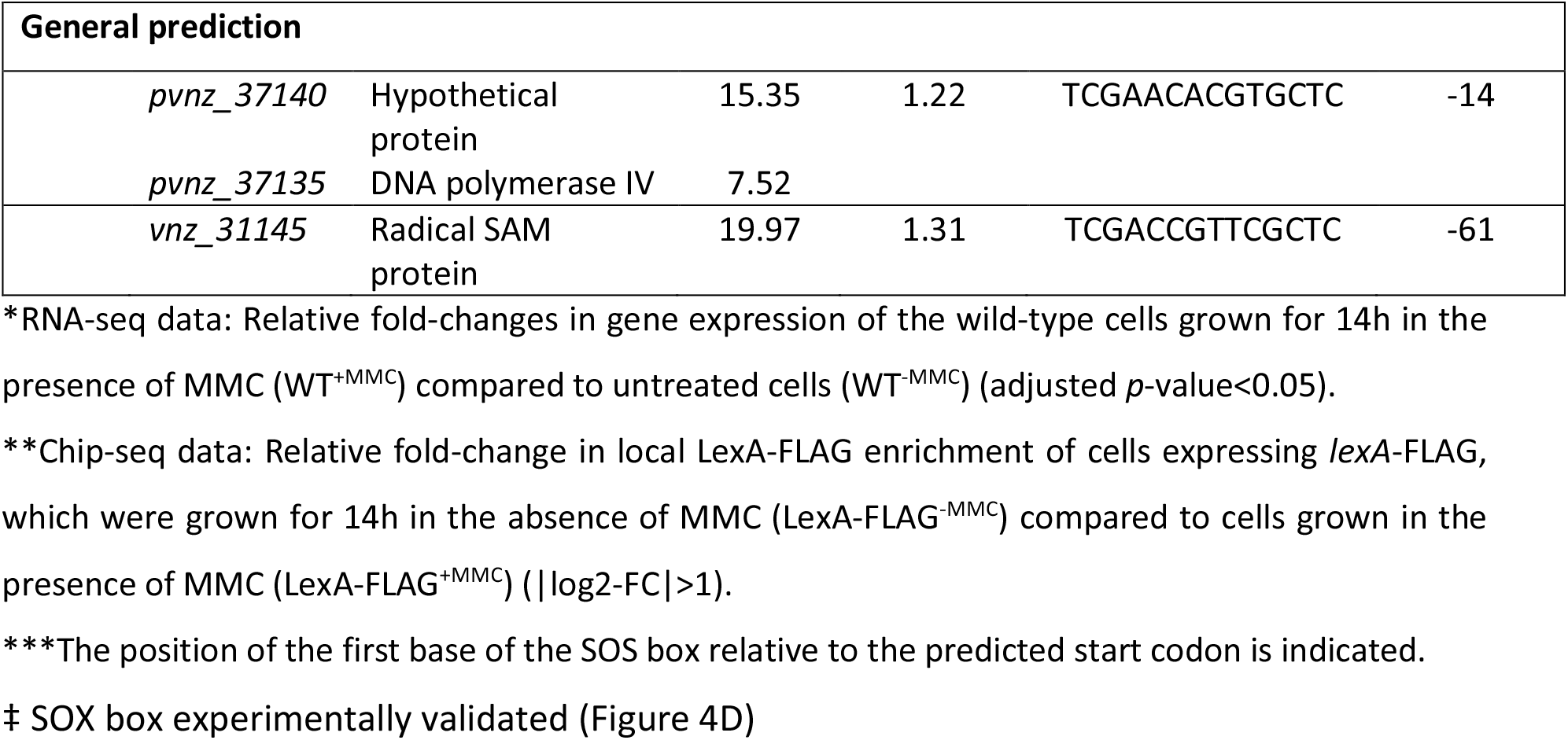
Selected members of the SOS regulon and co-regulated genes.

Moreover, the LexA regulon includes additional genes which could further contribute to the relief of DNA damage stress. These genes include *parE* (*vnz_27225*) which encodes an additional topoisomerase IV subunit; *vnz_31145* which shares similarity to radical SAM enzymes that repair DNA cross-links between thymine bases caused by UV radiation and an operon located on S. *venezuelae plasmid* pSVIJ1 consisting of a hypothetical protein and a error-prone DNA polymerase (*pvnz_37140-37135*).

### SOS boxes of DNA damage repair genes are tightly bound by LexA

Since the SOS-response involves the derepression of LexA target genes, we sought to study the effect of MMC-treatment on LexA binding *in vivo*. Therefore, and in parallel with our earlier experiments conducted under non-DNA damaging growth conditions, we performed ChIP-seq experiments following MMC-induced stress and subsequently compared LexA enrichment across the genome in untreated samples to ChIP-seq samples that had been obtained from cultures grown in the presence of 0.25 µg ml^-1^ MMC for 14h. In contrast to our untreated ChIP-seq experiments which demonstrated widespread binding of LexA across the genome, there was an observable decrease in genome-wide LexA binding in the presence of MMC (Figure 4A). This is in line with the observation that LexA protein levels are decreased under DNA-damaging conditions (Figure 2C). As in our earlier experiments, no significant enrichment was observed in the wild-type negative control under these conditions (Figure 4B). Of the 175 LexA binding sites determined here to be part of the SOS-response, MMC treatment resulted in reduced local enrichment of LexA at 112 genomic positions and the complete dissociation of LexA at 61 chromosomal loci. Both sets of loci also included the core DNA damage repair genes described above (Table 1 and Supplementary Figure 3A). Two chromosomal loci retained a similar level LexA enrichment under the conditions tested (Supplementary Table 4).

**Figure 4:**
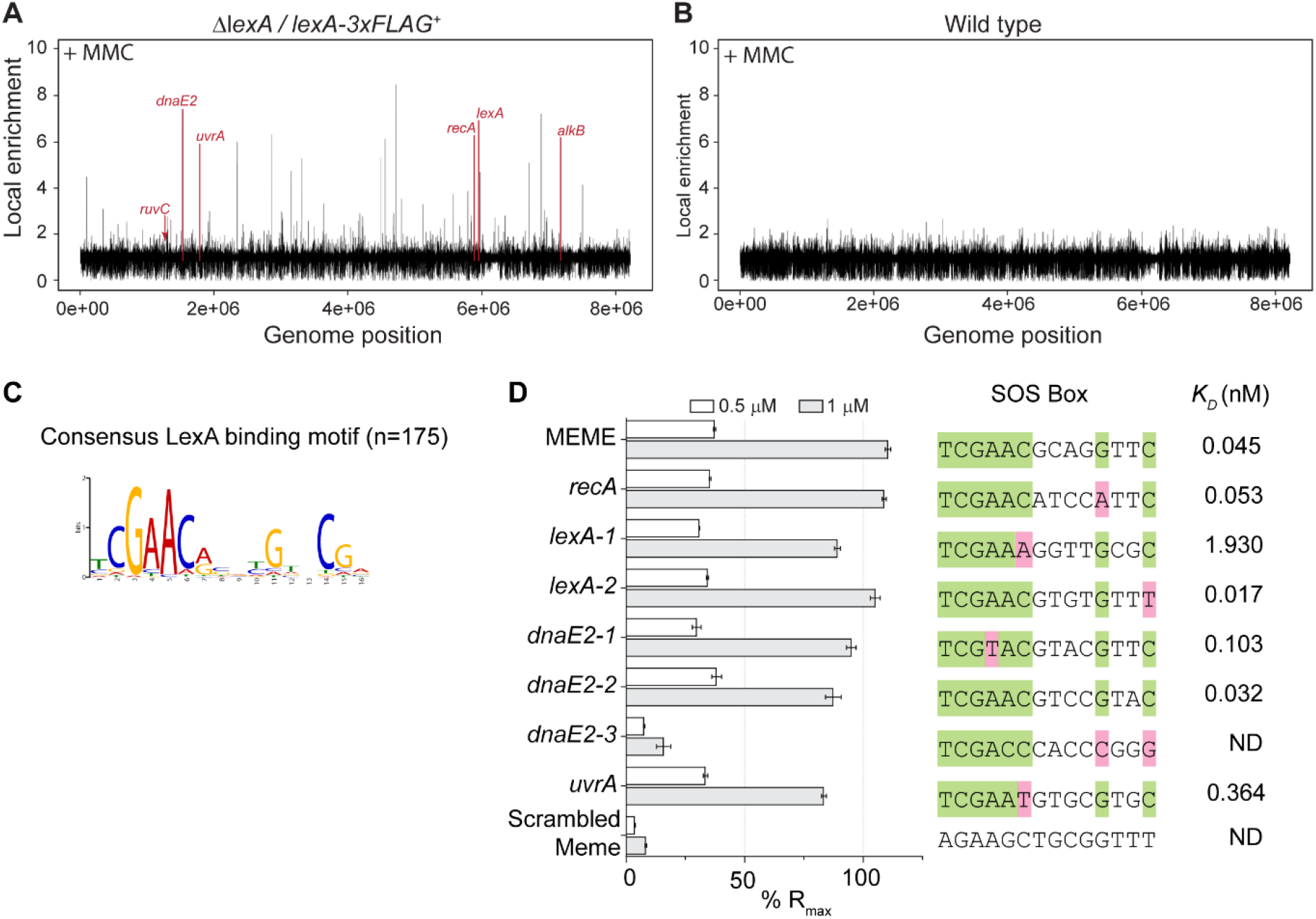
LexA binds tightly to SOS boxes of core DNA damage genes *in vitro*. Genome-wide ChIP peak distribution of the LexA regulon in the **(A)** Δ*lexA/lexA-FLAG*^*+*^ strain (KS74) and **(B)** the wild type. Strains were grown for 14h in the presence of 0.25 μg ml^-1^ MMC for 14h. Peaks of selected LexA regulon genes are labelled and coloured in red lines. **(C)** Identification of the LexA consensus motif (E-value 1.0e-252) by the MEME algorithm. The input of the analysis comprised ChIP-seq sequences of 100 nt length. The total number of sequences (n) is indicated. **(D)** Surface Plasmon Resonance (SPR) was used to determine the binding kinetics of LexA (0.5 μM and 1 μM) to 23-bp double-stranded DNA that contained individual target sequences located upstream of SOS genes involved in DNA repair. The target sequences (SOS box) used in these experiments are presented. Highly conserved nucleotides are shown in green, mismatches to the LexA consensus motif are shaded in magenta. A wider range of LexA concentrations was used to determine the binding affinity *K*_*D*_ of LexA to the individual target sequences (Supplementary Figure S3). The affinity for the *dnaE2-3* binding site and the scrambled MEME motif were not determined (ND) due to very little specific binding of LexA.

These findings prompted us to further examine the sequence-specificity of LexA binding. Using the motif-based sequence analysis tool MEME (21) and 100 nucleotide sequences located under the summit of each LexA ChIP-seq peak (n=175), we first identified the 16-bp consensus LexA DNA-binding motif tCGAAC-N_4_-GNNCGa (Figure 4C), which is an imperfect palindromic sequence, similar to previously reported LexA binding motifs from related and distant bacterial systems (3). We then compared this consensus LexA binding motif to the motifs generated from 100 nucleotide sequences associated with either ChIP-seq peaks that decreased (n=112) or disappeared (n=61) after MMC stress. This analysis did not reveal any significant differences in the sequence specificity of LexA binding *in vivo* (Supplementary figure 3B), indicating that additional factors might contribute to LexA enrichment and the differential response to MMC.

To further validate our *in vivo* ChIP-seq analysis, we examined the DNA binding specificity of purified *S. venezuelae* LexA using Surface Plasmon Resonance (SPR). 23-bp double-stranded oligonucleotides containing the consensus LexA DNA binding motif were tethered to a chip surface within an SPR flow cell. Purified (His)_6_-LexA was flowed over the test DNA and LexA binding was recorded by measuring the change in response units during LexA injection. Control experiments using double-stranded DNA with a scrambled LexA DNA binding sequence showed only a very weak association of LexA, supporting the idea that the observed binding was sequence-specific (Figure 4D). In addition, we performed SPR analysis using 23-bp duplex DNA that contained the putative LexA binding motifs identified upstream of *lexA, recA* and the two SOS genes *dnaE2* and *uvrA*. Based on our initial MEME analysis, we identified two potential LexA binding sites in the promoter region of *lexA* (*lexA1 and lexA2*) and three for *dnaE2 (dnaE2-1, dnaE2-2, dnaE2-3)*. In agreement with our ChIP-seq results, we observed clear binding of LexA to the predicted binding sites except for the *dnaE2-3* motif, suggesting that either this sequence may not represent a relevant LexA binding site or additional factor(s) are required for binding *in vivo* (Figure 4D). We repeated the SPR experiments, using sequentially increasing concentrations of LexA to enable the determination of the binding affinity (*K*_D_). These analyses showed a differential but overall strong binding of LexA to the tested duplex DNA, indicating that the induction of LexA target genes *in vivo* is under tight control in *Streptomyces* (Figure 4D and Supplementary Figure 3C). This may provide an explanation for the requirement of a prolonged exposure to genotoxic agents to induce the SOS response in *S. venezuelae* since LexA needs to first disassociate from its target sequence to become susceptible to RecA-stimulated self-cleavage (22). However, we cannot exclude the possibility that the overall stability of LexA also contributes to the delayed induction of the SOS response

## DISCUSSION

In this work, we confirm the presence of a functional DNA damage response and identify the core set of LexA-controlled SOS genes in *S. venezuelae*. While deletion of *recA* is not detrimental for most bacteria under non-DNA damaging growth conditions, deletion of *lexA* and the resulting constitutive activation of the SOS response is lethal in *E. coli* or causes cell filamentation in other bacterial species (23–26). *S. venezuelae* strains lacking either *recA* or *lexA* are viable, however, growth and sporulation are severely compromised in these mutant strains even under non-DNA damaging conditions. Available microarray data indicates that *recA* is constitutively expressed throughout the *S. venezuelae* life cycle (27). Given the importance of *recA* in genome maintenance (28), it is likely that the observed phenotypic defects in the *recA* mutant are a direct consequence of genome instability, which could result in large deletion in the chromosomal arms or defective chromosome segregation (8, 29). Furthermore, in the *lexA* null mutant permanent induction of the SOS response may result in the activation of a cell-cycle checkpoint that suppresses sporulation analogous to the SOS-dependent cell division inhibitors identified in diverse bacteria (30–32, 32, 33). Notably, *Streptomyces* lack clear homologs of these known cell division regulators, indicating the presence of a yet unidentified mechanism involved in coordinating DNA damage repair and cell division.

Our investigations revealed that the activation of the SOS response is delayed and requires a chronic exposure to DNA damaging agents in *S. venezuelae*. A similar slow activation of the SOS response were previously reported for other Streptomyces species and related Actinomycetota, showing that short treatments with DNA damaging agents were not sufficient to induce *recA* expression (34–36). The basis for this phenomenon is not fully understood but several factors may contribute to the delayed induction of the SOS response. For example, basal level of RecA is sufficiently high to repair DNA damage caused by short-term genotoxic stress without the need of inducing the SOS. It is also conceivable that LexA auto-cleavage is intrinsically slow compared to LexA from other organisms which is degraded within minutes (37, 38) or the co-protease function of RecA is inhibited by RecX, a negative regulator of RecA activity (36, 39). Although we cannot exclude the possibility that the intracellular accumulation of genotoxic agents is slowed down by increased efflux activity or properties of the *Streptomyces* cell envelope that limit the uptake of toxic compounds, these findings suggest that the regulation of the SOS response might be different in *Streptomyces*. Notably, the related Actinomycetota species *Mycobacterium smegmatis* employs two DNA damage response pathways, a RecA/LexA-dependent SOS response and a RecA/LexA-independent DNA damage response, with the latter one controlling the vast majority of DNA repair genes (40, 41). This alternative stress response involves a protein complex comprised of the two transcriptional regulators PafB and PafC, which bind to a DNA sequence motif that is distinct from the SOS box to activate the mycobacterial DNA damage response presumably upon binding single-stranded DNA (40–42). Although *Streptomyces* encode homologs of the PafBC system, an initial motif search suggested that the PafBC binding motif is not conserved in *S. venezuelae*. Thus, it remains to be shown whether a similar RecA/LexA-independent DNA response exists in *Streptomyces*.

Our results, however, establish the presence of a LexA-dependent DNA damage response in *Streptomyces*. In total, we identified 175 LexA binding sites that were associated with DNA-damage responsive genes that were both up and down-regulated following MMC-induced degradation of LexA. While the majority of these genes are of unknown function, about 36% of all differentially regulated genes are predicted to encode proteins involved in metabolism, signalling and other cellular processes. This clearly highlights the functional diversity of the gene categories that comprise the LexA regulon in *Streptomyces*. Importantly, our work identifies a core set of SOS-regulated DNA response genes involved in the regulation of the SOS response (*lexA, recA)*, DNA mutagenesis (*dnaE2-dinP*) and DNA repair (*ruvC-ruvA* operon and *alkB*) (3). We also identified several additional genes and previously unrecognized LexA target genes, involved in DNA decatenation (*parE*) (43, 44), UV-damage repair (*vnz_31145*) (45) and mutagenesis (*pvnz_37140-37135*) which could further contribute to restoring the integrity of the genome.

Furthermore, we detected a significant number of positively regulated LexA targets. Although we cannot exclude the possibility that the MMC-induced downregulation of LexA target genes might reflect the requirement for an additional transcriptional regulator(s) for induction, several examples of positive regulation by LexA have now been identified, (23, 46– 49). Two noticeable LexA targets in *S. venezuelae* that showed a significant downregulation upon MMC treatment include the structural genes for the developmental regulator WhiI and the small membrane protein SmeA which, together with its partner protein SffA, is involved in chromosome segregation during sporulation (18, 50). However, *whiI* and *smeA* expression is also controlled by additional developmental regulators, indicating the presence of a multi-layered regulatory network that controls the initiation of sporulation (16, 51). Together, our findings support the idea of a much more diverse cellular role of LexA in DNA damage stress resistance and *Streptomyces* development.

Our work further revealed that LexA displays a differential binding behaviour *in vivo*, indicating a graded response to DNA damage. These observations are in line with a hierarchical induction of LexA targets which is dependent upon both the level of DNA damage and the LexA dissociation rate constant at a given promoter (52–54). Indeed, our *in vitro* assays confirmed that in *S. venezuelae*, LexA binds tighter to the *recA, lexA2* and *dnaE2-2* operator sequences compared to the *lexA1/dnaE2-1* motif and the SOS box located in the promoter region of *uvrA*. This is likely to allow for a finely tuned response, balancing DNA damage surveillance, rapid repair of DNA lesions and mounting a full SOS response. Collectively, our work provides significant insight into the SOS response and how it might enable *Streptomyces* to survive in diverse environments.

## METHODS

### Bacterial strains and growth conditions

Strains, plasmids and oligonucleotides used in this work are listed in Supplementary Tables 5 and 6, respectively. *E. coli* strains were grown in LB or LB agar at 37°C supplemented with the following antibiotics when necessary: 100 μg ml^-1^ carbenicillin 50 μg ml^-1^ kanamycin, 25 μg ml^-1^ hygromycin, 50 μg ml^-1^ apramycin, or 25 μg ml^-1^ chloramphenicol. *S. venezuelae* NRRL B-6544 was grown in maltose-yeast extract-malt extract medium (MYM), composed of 50% tap water and 50% reverse osmosis water, and supplemented with R2 trace element (TE) solution at a ratio of 1:500. Liquid cultures were grown under aeration at 30 °C and at 250 rpm. MYM agar was supplemented with the following antibiotics when required: 5 μg ml^-1^ kanamycin, 25 μg ml^-1^ hygromycin, or 50 μg ml^-1^ apramycin.

### Construction and complementation of the *recA* null mutant

The Δ*recA* mutant strain (KS3) was generated using ‘Redirect’ PCR targeting (Gust et al., 2003 and 2004). The *recA* coding sequence (*vnz_26845*) on the cosmid vector Pl1-B2 (http://strepdb.streptomyces.org.uk/) was replaced by an *oriT-*containing apramycin resistance cassette, which was amplified from pIJ773 using the primer pair ks6 and ks7. The resulting disrupted cosmid (pKS100) was introduced into *S. venezuelae* by conjugation via *E. coli* ET12567/pUZ8002. Resulting exconjugants were screened for double crossover events (Apr^R^, Kan^S^) and deletion of *recA* was verified by PCR analysis using the primers ks8, ks9 and ks10. A representative *recA* null mutant was designated KS3. For complementation, plasmid pKS2 (*P*_*recA*_*-recA*) was introduced into the Δ*recA* mutant by conjugation.

Genomic DNA for genome sequencing of the ΔrecA mutant (KS3) and the parental wild-type strain was performed using the FastDNA kit for soil (MP Biomedicals Germany). Genomic DNA of biological replicate samples per strain was sequenced by MiGS (Pittsburgh, USA).

### Construction and complementation of the *lexA* null mutant

To generate the *ΔlexA* mutant strain (KS44), a plasmid carrying a second copy of *lexA* (pKS1) was first integrated at the phage BT1 attachment site of wild-type *S. venezuelae*. In the resulting strain (KS9), the native *lexA* coding sequence *(vnz_27115*) was then replaced by the *apr-oriT* cassette using the REDIRECT PCR targeting technology and cosmid Pl2-B2 as described above, using the primer pair ks1 and ks2 to PCR amplify the apr-oriT cassette from pIJ773. The mutagenized cosmid (pKS200) was introduced into KS9 by conjugation via ET12567/pUZ8002 and exconjugants that had successfully undergone double crossover events were identified by screening for apramycin-resistance and kanamycin sensitivity yielding strain KS25. The Δ*lexA::apr* allele of KS25 was then moved back into wild-type *S. venezuelae* via generalised SV1 phage transduction as described by Tschowri et al. (55). Transduction of the Δ*lexA::apr* allele was confirmed by PCR analysis using the primers ks3, ks5, and ks32. A representative Δ*lexA* null mutant strain was designated KS44. For complementation, plasmid pKS1 (*P*_*lexA*_*-lexA*) or pKS3 (*P*_*lexA*_*-lexA-3xFLAG*) was introduced into the Δ*lexA* mutant by conjugation, yielding strain KS57 and KS74, respectively.

### RNA preparation, sequencing (RNA-seq) and analysis

RNA-seq was conducted in biological triplicate using the protocol described by Bush et al. (56). Triplicate cultures of wild-type *S. venezuelae* were grown in 30 ml MYM cultures with shaking (250 rpm) for 14h at 30°C, “treated” cultures were grown with the addition of 0.25 µg ml^-1^ mitomycin C (MMC) (Merck Life Sciences UK Ltd). Mycelial pellets from untreated and treated cultures were washed in 1x PBS before lysis and RNA purification using the RNEasy Kit (Qiagen). Purified RNA samples were treated with on-column DNase I (Qiagen), followed by an additional DNase I treatment (Turbo DNA-free, Invitrogen) and the absence of DNA contamination was confirmed by PCR amplification of the house-keeping gene *hrdB* (*vnz_27210*) using the primer pair mb1 and mb2. Library construction, rRNA depletion and paired-end Illumina Hiseq sequencing (2×150 bp configuration) was performed by Genewiz (NJ, USA).

The quality of the obtained sequencing reads was checked using FastQC (https://www.bioinformatics.babraham.ac.uk/projects/fastqc/) and the reads were mapped to the genome of *S. venezuelae* NRRL B-65442 (NCBI Reference Sequence: NZ_CP018074.1) using the Bowtie2 alignment tool (57) and subsequently sorted and indexed using the Samtools package (58). A custom Perl script was used to make a saf file for genes in the *S. venezuelae* genome. The featureCounts tool of the BioConductor package “Rsubread” was used to count the reads mapping to every gene in the *S. venezuelae* genome. The counts were read into a DGEList object of the edgeR package of R and a quasi-likelihood negative binomial generalized log-linear model was fitted to the data using the glmQLFit function of edgeR. Genewise statistical tests were conducted using the glmQLFTest function of edgeR (59). The “topTags” function was used to arrive at a table of differentially expressed genes (59). Genes were classed as differentially expressed genes (DEGs) if the log2-fold-change (treated/untreated) was greater than or equal to 1, or less than or equal to -1 and had a false discovery rate (FDR, adjusted *p*-value) of less than or equal to 0.05.

### Chromatin immunoprecipitation-sequencing (ChIP-seq) and analysis

ChIP-seq was performed in biological replicates as described previously (16) using a *S. venezuelae* Δ*lexA* strain that was complemented with a *lexA-3xFLAG* fusion (KS74). Cells were grown in MYM medium in the absence (untreated) and presence of 0.25 µg ml^-1^ MMC for 14h (14-MMC) at 30°C. Library construction and sequencing were performed by Genewiz (NJ, USA) using Illumina Hiseq (2 × 150bp configuration, trimmed to 100bp). Obtained sequencing reads were aligned to the *S. venezuelae* NRRL B-65442 genome (NCBI Reference Sequence: NZ_CP018074.1) using the Bowtie 2 alignment tool (57). The resultant bam files were sorted and indexed using the Samtools package (58). A custom Perl script was used to make a saf file for every 30-nucleotide section in the *S. venezuelae* genome with adjacent sections overlapping by 15 nucleotides. The featureCounts tool of the R package Rsubread was used to count the number of reads mapping to every 30-nucleotide section in the saf file made above. A custom R script was then employed to calculate a local enrichment for every 30-nucleotide section (overlapping by 15 nucleotides) by comparing the read density in the section to the read density in the 3000 nucleotides around the section. The local enrichment in the corresponding sections of the controls were subtracted from the local enrichment of the ChIP samples. Custom Perl and R scripts were used to list out genes to the left, right, and overlapping genome sections with a local enrichment of 2 or more. In addition, genes had to be in the right orientation and within 300 nucleotides of the enriched region for association with LexA-3xFLAG binding sites. Intragenic LexA bindings sites (local enrichment of >2) were defined as genomic positions that were located more than 100 bp downstream of the annotated start codon of a gene and which were not considered to be part of a promoter region of the downstream gene. Of the 208 intragenic LexA-3xFLAG binding sites, 176 binding sites were detected in proximity of a transcription start site (based on the “14h” TSS dataset publicly available at EMB-EBI E-MTAB-10690) (11).

To assess the effect of MMC, the enrichment of the 175 LexA-3xFLAG ChIP-seq peaks located upstream of transcriptional units was assessed based on the differential response to MMC stress (adjusted *p*-value *p*<0.05 and fold change cut-off |log2-fold change|>1) and on the following criteria: (i) Complete dissociation of LexA-3xFLAG from a binding site upon MMC-treatment was judged to have occurred when the enrichment value (log2-fold-change) in the MMC-treated ChIP-seq dataset (LexA-3xFLAG-14hMMC/WT-14hMMC) was less than 0.25. (ii) No dissociation of LexA-3xFLAG from a binding site upon MMC-treatment was called when the enrichment value (log2-fold-change) calculated from the comparison of the treated and untreated ChIP-seq datasets (LexA-3xFLAG-14hMMC/LexA-3xFLAG-untreated) was less than 0.2. (iii) Partial dissociation of LexA-3xFLAG from a binding site upon MMC-treatment was defined when peaks displayed a local enrichment value (log2-fold-change) in the MMC-treated ChIP-seq dataset (LexA-3xFLAG-14hMMC/WT-14hMMC) of more than 0.25 and the enrichment value (log2-fold change) calculated from the comparison of the treated and untreated ChIP-seq datasets (LexA-3xFLAG-14hMMC/LexA-3xFLAG-untreated) was greater than 0.2.

### Functional analysis of LexA regulon

Analysis for the enrichment of functional classes was performed on proteins encoded by the 176 genes that were associated with intragenic LexA binding sites and the 175 differentially expressed genes that are subject to LexA regulation (Supplementary Table 2 and 4). To predict conserved protein features and to assign COG identifiers (clusters of orthologous groups), the queries were searched against the Position Specific Scoring Matrix (PSSM) database (NCBI) using the rpsblast program (60). Where more than one COG identifier was reported, genes were inspected for their genetic context and predicted function and were manually assigned a COG class.

### Motif prediction from LexA-enrichment sites

100 nucleotide sequences surrounding the centre position of each of the 175 enrichment sites determined by our ChIP-analysis were used as input for the MEME suite (21) (Supplementary Table 4), with an *S. venezuelae* background model, to search for a conserved LexA binding motif (SOS box). Site distribution was set to “Zero” or “One Occurrence Per Sequence”, and the MEME algorithm was set to search for three motifs on both strands with a width of 6-500 nucleotides.

### Automated Western blotting (WES)

For determining LexA and LexA-FLAG levels, the Δ*lexA*/*lexA-3xFLAG* strain (KS74), and the wild type were used. To confirm specific binding of the antibodies, the wild type or the Δ*lexA* deletion strain (KS44) were used as the respective negative control. Duplicate cultures of each strain were grown in liquid MYM medium for 14h without (untreated) and with 0.25 µg ml^-1^ MMC, 3 µg ml^-1^ ciprofloxacin, or 3 mM MMS. After 14h, untreated cultures were exposed for 1 and 2h to 0.25 µg ml^-1^ or 2.5 µg ml^-1^ MMC, 3 µg ml^-1^ ciprofloxacin, and 5 mM MMS, respectively. At the end of the incubation time, 2-5 ml of each MYM culture was removed and washed once with 1x PBS. Samples of mycelium were resuspended in 0.4 ml ice-cold sonication buffer [20 mM Tris pH 8.0, 5 mM EDTA, 1x EDTA-free protease inhibitors (Sigma Aldrich)] and sonicated (5x 15 sec on/15 sec off) at 4.5-micron amplitude. Lysates were then centrifuged at 16,000 x g for 15 min at 4°C to remove cell debris. Total protein concentration was determined using the Bradford assay (Biorad). To detect LexA-FLAG or LexA abundance in cell lysates, 2 µg of total protein for each sample was loaded in duplicate into a microplate (ProteinSimple #043-165) together with either anti-FLAG antibody (Sigma F4725) diluted 1:100 or polyclonal anti-LexA antibody diluted 1:200. LexA-FLAG and LexA levels, were then assayed using the automated Western blotting machine (ProteinSimple, San Jose, CA), according to the manufacturer’s guidelines.

Virtual Western blots related to Figure 2C and Supplementary Figure 2C were generated using the Compass software for simple western (Version 6.0.0) and uncropped images of representative blots are shown in Supplementary Figure S4.

### Fluorescence microscopy

To visualize the expression of the *P*_*recA*_*-mcherry* fusion in *S. venezuelae* (KS80), cells were grown in triplicate in liquid MYM medium with and without MMC, ciprofloxacin and MMS as described. Cells obtained from each culture were immobilized on a 1% agarose pad and visualized using a Zeiss Axio Observer Z.1 inverted epifluorescence microscope fitted with a sCMOS camera (Hamamatsu Orca FLASH 4), a Zeiss Colibri 7 LED light source, a Hamamatsu Orca Flash 4.0v3 sCMOS camera. Images were acquired using a Zeiss Alpha Plan-Apo 100x/1.46 Oil DIC M27 objective with an excitation/emission bandwidth of 577-603 nm/614-659 nm to detect fluorescence emitted by the mCherry reporter fusion. Still images were collected using the Zen Blue software and further analyzed using the Fiji imaging software (61). To normalise the mCherry fluorescence intensity across different MMC treatments, images were corrected for background fluorescence.

### Cryo-scanning electron microscopy

*Streptomyces* samples were mounted on an aluminium stub using Tissue Tek^R^ (Agar Scientific Ltd, Stansted, England) and plunged into liquid nitrogen slush. The sample was transferred onto the cryo-stage of an ALTO 2500 cryo-system (Gatan, Oxford, England) attached to an FEI Nova NanoSEM 450 field emission SEM (Thermo Fisher Scientific, Eindhoven, The Netherlands). Sublimation of surface frost was performed at -95°C for four minutes before sputter coating with platinum for 150 sec at 10 mA. The sample was moved onto the main cryo-stage in the microscope, held at -125°C, and imaged at 3 kV, spot 3, and digital TIFF files were stored.

### Growth assays

To determine viability of *S. venezuelae* strains on solid medium 10e^5^ colony forming units per ml^-1^ (spores or mycelia fragments) were used to prepare a 10-fold serial dilution series and 3 μl of each dilution series was subsequently spotted onto MYM agar containing either no antibiotic, 0.25 µg ml^-1^ MMC, 3 µg ml^-1^ ciprofloxacin, or 3 mM MMS. Plates, in biological triplicate, were incubated at 30°C for 3-5 days and subsequently imaged.

To assay the growth of *S. venezuelae* in liquid MYM, OD_600_ measurements were made using a plate reader (SPECTROstar Nano by BMG Labtech) set at 30°C and shaking at 700 rpm. 800 µl of MYM supplemented with increasing concentrations of MMC, ciprofloxacin, or MMS was loaded into the wells of a 48-well-plate (677102 Greiner Bio-One) and inoculated with either 5 µl spores or 10 µl of mycelium. OD_600_ measurements were recorded every 15 min for 24 hours and data was subsequently visualized using Graphpad Prism (Version 9.3.1). Each experiment was performed in triplicate.

### Purification of His_6_-LexA

To purify His_6_-LexA, *E. coli* NiCo21 (DE3) cells were transformed with the pLysS and the pKS16 plasmid. Cells were grown at 30°C in LB medium containing 25 µg ml^-1^ chloramphenicol and 100 µg ml^-1^ carbenicillin until OD_600_ 0.5 before the addition of 0.5 mM IPTG (isopropyl β -D-1-thiogalactopyranoside) to induce protein production. Cultures were incubated with shaking at 30°C for 4h and then harvested by centrifugation. Cell pellets were resuspended in 50mM Tris pH 10.5, 1 M NaCl, 10% glycerol with protease inhibitor (EDTA-free, Merck) and lysed by sonication for 14 cycles at 18-micron amplitude, 15 sec on/ 10 sec off. Lysates were centrifuged at 26,000 x g for 45 min at 4°C to remove cell debris and passed through a 0.45 µm filter before loading on a HisTrap column via the ÄKTA pure system. (GE Healthcare). His_6_-LexA was then eluted using an increasing concentration of imidazole. Fractions containing His_6_-LexA were pooled and subjected to size exclusion chromatography on a HiLoad 16/600 Superdex 200 pg column (GE Healthcare) in 50mM Tris pH 10.5, 200 mM NaCl, 10% glycerol, 0.5 mM DTT. The concentration of pooled protein fractions was determined by Bradford assay and His_6_-LexA was subsequently stored at -80°C until further use. To produce antibodies against LexA from *Streptomyces*, His_6_-LexA was overexpressed and purified as described above, and a total amount of 3.54 mg of purified protein was sent to Cambridge Research Biochemicals (UK) to be used to raise antibodies in rabbits.

### Surface Plasmon Resonance (SPR)

All SPR experiments were performed using a Biacore 8K instrument (Cytiva). All measurements were performed with a single sensor chip SA (Cytiva), using the ReDCaT system as described previously (62). Complementary single-stranded oligonucleotides containing the individual LexA binding sequences, or a scrambled binding motif were annealed and subsequently diluted to a working concentration of 1 µM in HPS-EP+ buffer (0.01 M HEPES pH 7.4, 0.15 M NaCl, 3 mM EDTA, 0.005% v/v Surfactant P20). Each DNA was designed to contain the sequence of interest, and a single stranded extension on the reverse primer which anneals to the ReDCaT sequence on the chip, enabling DNA of interest to be bound and stripped after each experiment. For each experiment double-stranded DNA (with 20 base single-strand extension) was flowed over one flow cell on the chip at a flow rate of 10 µl min^-1^ for 60 sec to allow annealing to the ReDCaT linker via the complementary DNA. Purified *S. venezuelae* His_6_-LexA (diluted in HBS-EP+ buffer) was flowed over both the blank cell surface and the one containing the DNA to ensure the response seen was specific to binding to the DNA. To assess whether LexA bound the DNA, 1 µM and 0.5 µM His_6_-LexA was used at a flow rate of 50 µl min^-1^ for 120 sec before switching back to buffer flow to allow dissociation for 120 sec. The DNA and any remaining protein were then removed with a wash over both flow cells at 10 µL min^-1^ with 1 M NaCl, 50 mM NaOH. DNA sequences were tested in triplicate at two different protein concentrations and the amount of binding was recorded as response units (RUs).

To determine LexA binding kinetics, a single cycle kinetics approach was used. DNA was initially captured as previously described on the test flow cell and then increasing concentrations of His_6_-LexA (0.78125 nM, 1.5625 nM, 3.125 nM, 6.25 nM, 12.5 nM, and 25 nM) were injected over both the blank and DNA bound flow cells at a flow rate of 100 µl min^-1^ for 120 sec. At the end of all the His_6_-LexA injections, HPS-EP+ buffer was flowed over the surface for 1200 sec to measure the dissociation. The DNA and any remaining bound protein were then removed using the 1M NaCl and 50 mM NaOH wash. The recorded sensorgrams were analyzed using Biacore Insight Evaluation software version 3.0.11.15423 (Cytiva). The following formulae were used to calculate R_max_ and %R_max_-values, respectively: Rmax=(Mwt LexA)/(Mwt DNA) × RU × n × 0.78 and %R_max_= (Analyte binding RU)/R_max_ × 100. Where Mwt is the molar mass, RU is the DNA capture response, 0.78 is a constant used for estimating responses for DNA/protein interactions and n is the binding stoichiometry. Raw data used to calculate R_max_ and %R_max_ is shown in Supplementary Table 6. To calculate LexA binding kinetics, a fit was applied using a predefined kinetics 1 to 1 binding model provided with the Biacore 8K Evaluation Software 3.0.11.15423 (Cytiva). Data was plotted using Graphpad Prism (Version 9.3.1).

## Data summary

The authors confirm that all supporting data have been provided within the article or through supplementary data files. Raw RNA-seq and ChIP-seq data have been deposited in the MIAME-compliant ArrayExpress database (https://www.ebi.ac.uk/arrayexpress/) under accession number E-MTAB-11556 and E-MTAB-11603.

## Acknowledgments

This work was funded by a Royal Society University Research Fellowship (URF\R1\180075) to SS and by the BBSRC Institute Strategic Program grant BB/J004561/1 to the John Innes Centre. KJS’s PhD studentship was funded by the Royal Society Enhancement Award (RGF\EA\181026).

