## Supplementary figures for "Genome-wide identification of the LexA-mediated DNA damage response in *Streptomyces venezuelae*"

SUPPLEMENTARY INFORMATION

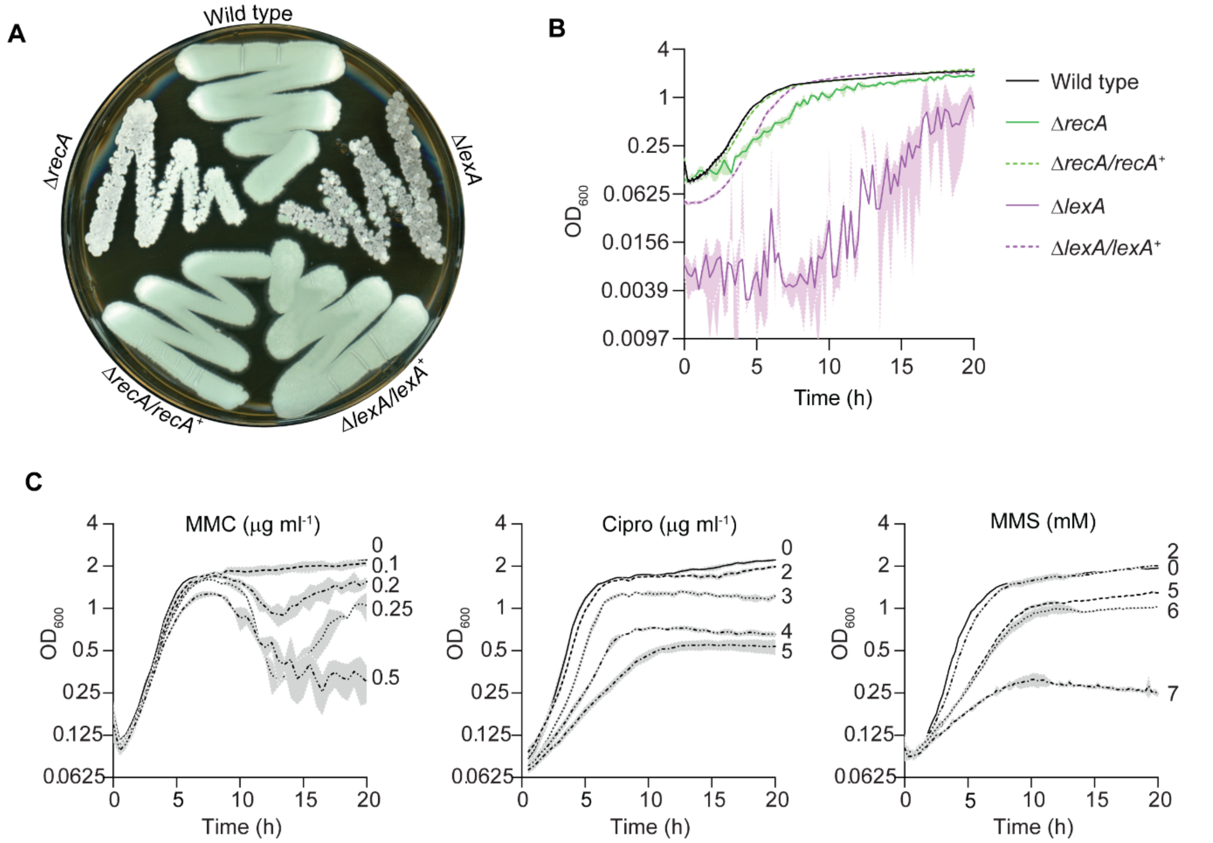


**Supplementary Figure 1: Deletion of *lexA* and *recA* and the treatment with genotoxic agents impair normal growth of *S. venezuelae*. (A)** Shown are the phenotypes of wild-type *S. venezuelae*, the constructed Δ*recA* (KS3) and Δ*lexA* (KS44) null mutants and the complemented strains Δ*recA/recA*+ (KS14) and Δ*lexA/lexA*+ (KS57). Strains were grown on MYM agar and imaged after 4 days. **(B)** Growth curves in liquid MYM for the wild type, the Δ*recA* (KS3) and Δ*lexA* (KS44) null mutants and the complemented strains Δ*recA/recA*+ and Δ*lexA/lexA*+ (KS14, KS57). **(C)** Growth curves in liquid MYM for the wild type in the absence and presence of increasing concentrations of mitomycin C (MMC), ciprofloxacin (Cipro) and methyl methanesulfonate (MMS). Growth curves shown in (B) and (C) depict the mean growth rate and the standard error obtained from biological triplicate experiments.

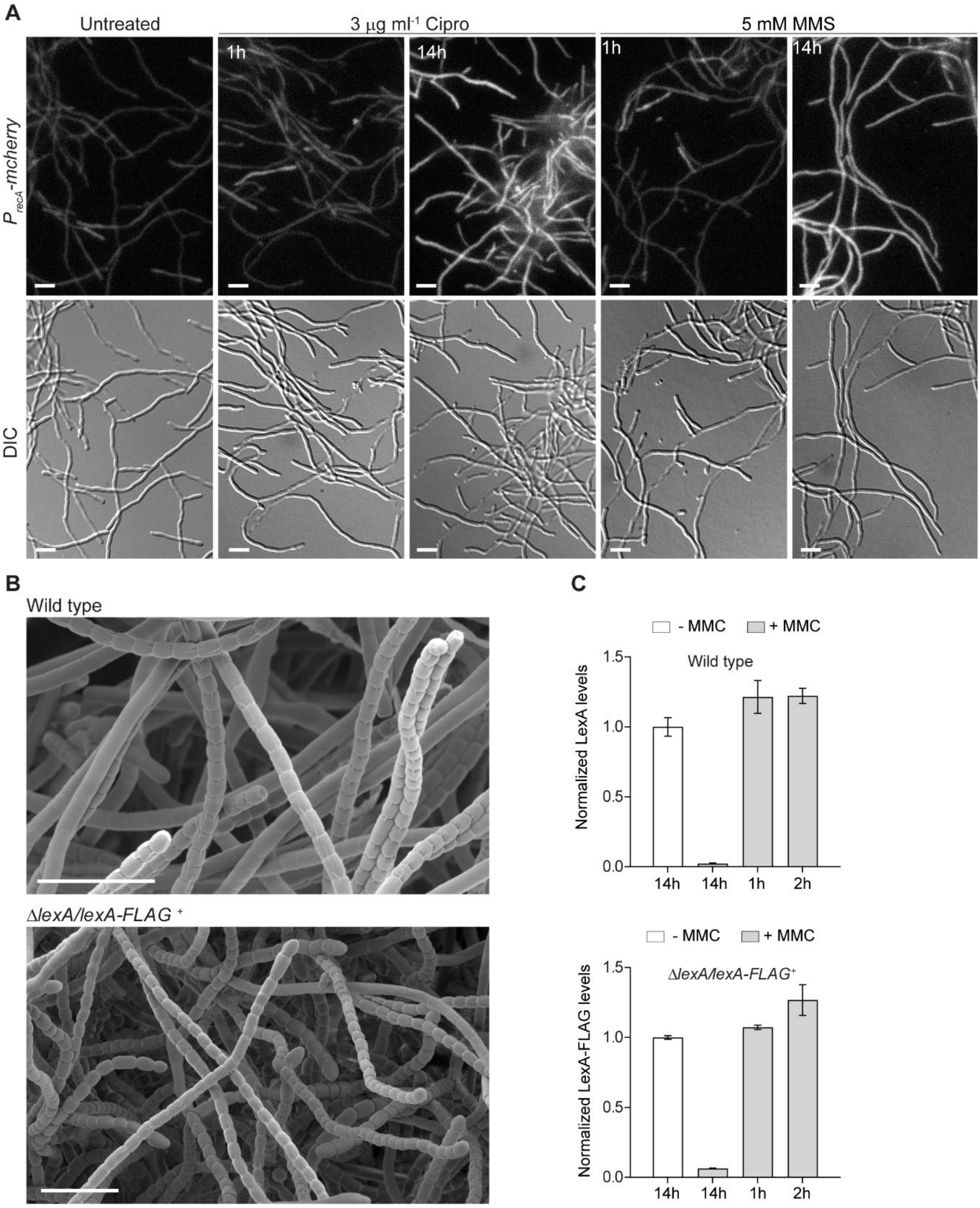


**Supplementary Figure 2: Induction of the SOS response requires extended exposure to genotoxic agents. (A)** Fluorescence micrographs and corresponding differential interference contrast images (DIC) of wild-type *S. venezuelae* expressing a *recA-mcherry* promoter gene fusion ($P_{recA}$*-mcherry,* KS80). Cells were grown in the absence or presence of 3 μg ml$^{-1}$ ciprofloxacin (Cipro) and 5 mM methyl methanesulfonate (MMS) for 1h or 14h. Shown are representative images from biological triplicate experiments. Scale bars: 5 μm. **(B)** Representative cryo-scanning electron micrographs showing sporulating aerial hyphae of the wild type and the complemented Δ*lexA* null mutant expressing a l*exA*-*FLAG* gene fusion *in trans* (KS74). Scale bars: 5 μm **(C)** Automated Western blot analysis showing the LexA protein stability in the wild type compared to LexA-FLAG in the complemented Δ*lexA/lexA-FLAG*$^{+}$ strain (KS74). Both strains were treated with 0.25 μg ml$^{-1}$ mitomycin C (MMC) for 1h, 2h and 14h. LexA and LexA-FLAG abundance was detected using a polyclonal α-LexA antibody and a monoclonal α-FLAG antibody, respectively. Protein levels were normalized to the untreated control. Shown are the mean protein levels and the standard error obtained from two biological replicate experiments. Representative virtual Western blots are shown in Supplementary Figure 4B.

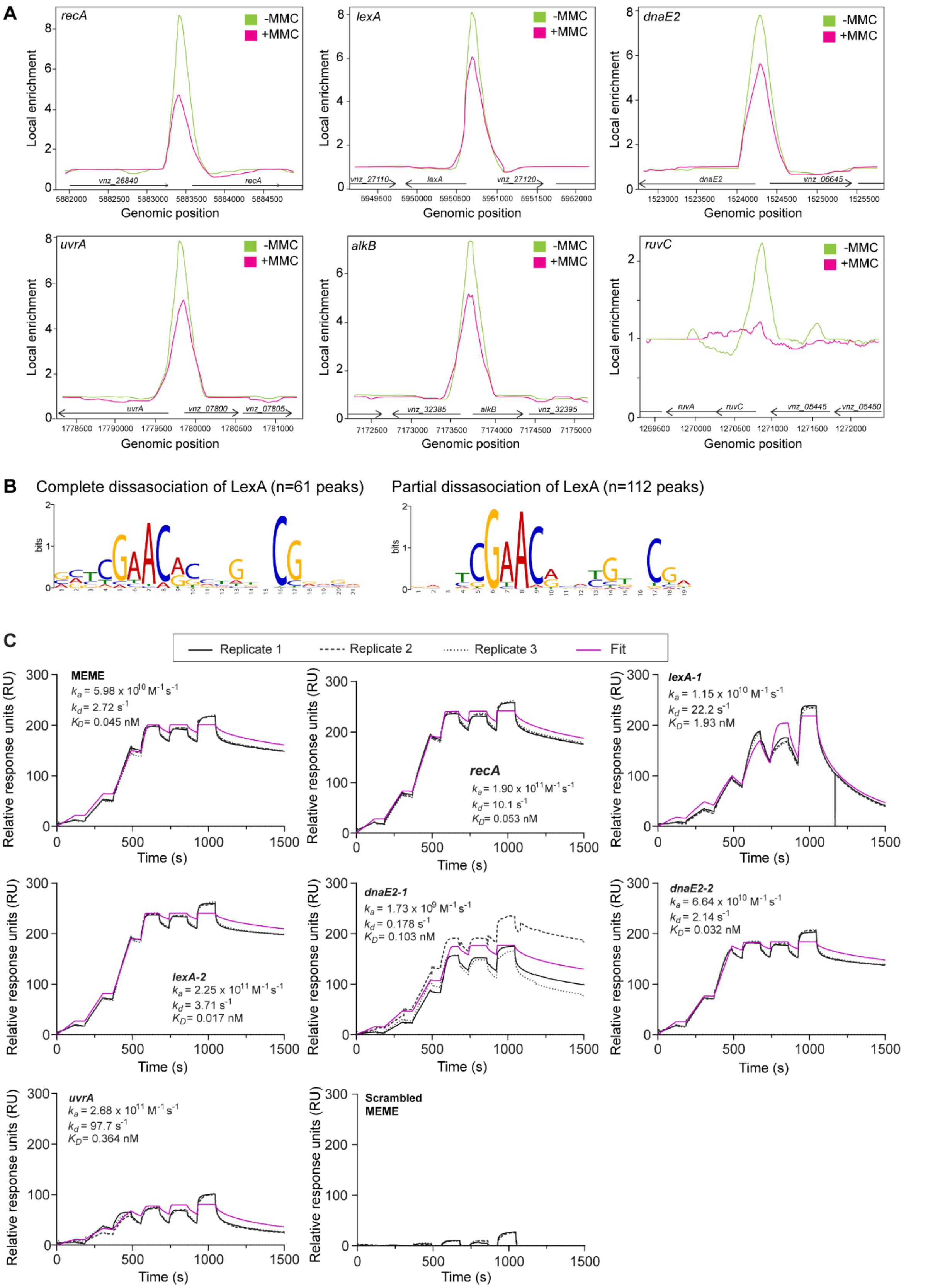


**Supplementary Figure 3: LexA binding to DNA-damage responsive target sequences *in vivo* and *in vitro*. (A)** ChIP-seq data for LexA binding to the promoter region of the DNA-damage inducible genes *recA*, *lexA*, *dnaE2*, *uvrA*, *alkB* and *ruvC*. Shown is the local enrichment of LexA following the growth of the strain Δ*lexA/lexA-FLAG*$^{+}$ (KS74) in the absence (green) and in the presence (magenta) of 0.25 μg ml$^{-1}$ mitomycin C (MMC) for 14h. Arrows and vnz-numbers indicate orientation and genomic identifier of surrounding genes, respectively. **(B)** Identification of the LexA binding motifs associated with ChIP-seq data showing the complete dissociation (left, E-value 1.5e-61) or a reduced enrichment (right, E-value 8.4e-177) of LexA in response to MMC stress. Sequence logos were determined by the MEME algorithm using ChIP-seq sequences of 100 nt length. The total number of sequences (n) is indicated. **(B)** Determination of the LexA binding kinetics for selected DNA binding motifs upstream of selected SOS genes using single-cycle kinetics SPR with sequentially increasing concentrations of LexA (0.78125, 1.5625, 3.125, 6.25, 12.5, and 25 nM). Binding of LexA to the target sequences was recorded in triplicate and expressed as response units (RU). A predefined kinetics 1 to 1 binding model provided with the Biacore 8K Evaluation Software was used to generate the fit and calculate the LexA binding kinetics.

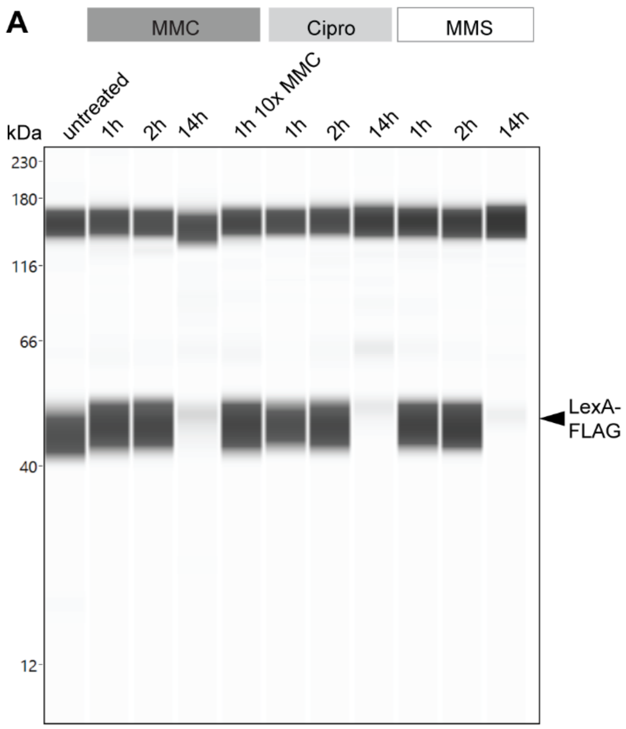


**Supplementary Figure 4: Virtual Western blots related to Figure 2c and Supplementary Figure 2C. (A)** Equal amounts of protein lysate obtained from strain KS74 in the absence or presence of MMC were probed with anti-FLAG antibody. **(B)** Analysis of LexA and LexA-FLAG abundance in the wild type and in KS74, respectively, following the treatment with MMC. Equal amounts of protein lysate were probed with either polyclonal anti-LexA or anti-FLAG antibody. Shown are representative virtual Western blots of biological duplicate experiments.

**Supplementary data provided as XLSX files**

**Supplementary Table 1:** *S. venezuelae* LexA regulon. ChIP-seq data obtained from *S. venezuelae* expressing a *lexA-3xFLAG* fusion (KS74) grown in MYM for 14h in the absence (untreated) and the presence of 0.25 μg ml-1 MMC (14h MMC). Only LexA enrichment sites that displayed a Log2-fold change>1 between the untreated and the MMC-treated samples and which are located within -200 to +100 bp relative to the start codon of the first gene in a transcriptional unit are listed. Experiments were performed in biological duplicate.

**Supplementary Table 2:** Intragenic LexA binding sites. ChIP-seq data obtained from S. venezuelae expressing a lexA-3xFLAG fusion (KS74) grown in MYM. Only LexA enrichment sites that displayed a Log2-fold change>1 between and which are located >100 bp downstream relative to the start codon of the first gene in a transcriptional unit and which were not part of previously defined promoter region are listed. Intragenic LexA binding sites associated with genes involved in DNA replication, repair and cell division are highligthed in grey.

**Supplementary Table 3:** Global response of *S. venezuelae* to DNA damage stress. RNA-seq data listing all genes that displayed a differential expression when comparing the wild type (WT) *S. venezuelae* grown in MYM for 14h with 0.25 μg $ml^{-1}$ MMC (+MMC) to untreated cells (-MMC). The following thresholds were applied: adjusted *p*-value $p<0.05$, |log2-fold change|>1. RNA-seq experiments were performed in biological triplicate.

**Supplementary Table 4**: *S. venezuelae* SOS regulon**.** Integrated RNA-seq and ChIP-seq data showing genes differentially expressed in response to MMC treatment and associated with a predicted LexA binding site. Only differentially expressed genes with an adjusted *p*-value $p<0.05$ and a |log2-fold change|>1 that had a predicted LexA binding site -200 to +100 bp relative to the start codon were considered.

**Supplementary Table 7:** Supplementary Table 6: Raw SPR used to calculate Rmax and %Rmax.

**Supplementary Table 5:** Strains and plasmids used in this study.

| Strain | Description | Construction | Source |
|---|---|---|---|
| *Escherichia coli* | | | |
| TOP10 | *F– mcrA Δ(mrr-hsdRMS-mcrBC) Φ80lacZΔM15 ΔlacX74 recA1 araD139 Δ(ara leu) 7697 galU galK rpsL (StrR ) endA1 nupG* | Cloning | Invitrogen |
| ET12567/pUZ8002 | *F– dam13::Tn9 dcm6 hsdM hsdR recF143:: Tn10 galK2 galT22 ara-14 lacY1 xyl-5 leuB6 thi-1 tonA31 rpsL hisG4 tsx-78 mtl-1 glnV44* | ET12567 with helper plasmid pUZ8002 | (1) |
| BW25113/pIJ790 | *Δ(araD-araB)567 ΔlacZ4787(::rrnB-4) lacIp-4000(lacIQ), λ–rpoS369(Am) rph-1 Δ(rhaD-rhaB)568 hsdR514* | BW25113 containing λ RED recombination plasmid pIJ790 | (2) |
| DH5α | *F- endA1 glnV44 thi-1 recA1 relA1 gyrA96 deoR nupG Φ80dlacZΔM15 Δ(lacZYA-argF)U169, hsdR17(rK - mK + ), λ–* | Cloning | New England Biolabs |
| NiCo21 (DE3) pLysS | *can::CBD fhuA2 [lon] ompT gal (λ DE3) [dcm] arnA::CBD slyD::CBD glmS6Ala ΔhsdS λ DE3 = λ sBamHIo ΔEcoRI-B int::(lacI::PlacUV5::T7 gene1) i21 Δnin5; pLysS: p15A replicon, T7 lysozyme coding sequence,* $Cm^R$ | Protein expression | New England Biolabs |
| *Streptomyces venezuelae* | | | |
| NRRL B-65442 | *Wild Type (WT)* | | (3) |
| KS3 | *ΔrecA::apr* | chromosomal *recA* locus deleted using pKS100 | This study |
| KS9 | WT *attB*$_{\phi BT1}$*::lexA* | pKS1 integrated at ϕBT1 attachment site | This study |
| KS14 | *ΔrecA::apr attB*$_{\phi BT1}$*::recA* | pKS2 integrated at ϕBT1 attachment site of KS3 | This study |
| KS18 | *ΔrecA::apr attB*$_{\phi BT1}$*::pIJ10770* | pIJ10770 integrated at ϕBT1 attachment site of KS3 | This study |
| KS25 | *ΔlexA::apr attB*$_{\phi BT1}$*::lexA* | Chromosomal *lexA* locus in KS9 deleted using pKS200 | This study |
| KS44 | *ΔlexA::apr* | *ΔlexA::apr* allele from KS25 transduced into the WT using SV1 generalised phage transduction | This study |
| KS57 | *ΔlexA::apr attB*$_{\phi BT1}$*::lexA* | pKS1 integrated at ϕBT1 attachment site of KS44. | This study |

| | | | |
|---|---|---|---|
| KS74 | *ΔlexA::apr* $attB_{\phi BT1}$*::lexA-3xFLAG* | pKS3 integrated at ϕBT1 attachment site of KS44. | This study |
| KS80 | WT $attB_{\phi BT1}::{}_{p}$*recA-mCherry* | pKS22 integrated at ϕBT1 attachment site | This study |
| Plasmids | | | |
| pIJ773 | pBluescript KS (+) containing the apramycin resistance gene *apr* and *oriT* of plasmid RP4, flanked by FRT sites (AprR). Used as template for the amplification of the *apr oriT* cassette for 'Redirect' PCR targeting, $Apr^R$_ | | (4) |
| pIJ10770 | Cloning vector for the conjugal transfer of DNA from *E. coli* to *Streptomyces* spp. Integrates at the ΦBT1 attachment site. $Hyg^R$ | | (5) |
| pIJ10772 | Modified pIJ10770, carries *mcherry* for construction of C-terminal fluorescent gene fusion, $Hyg^R$ | | (5) |
| pET15b | Expression vector. $Kan^R$ | | Novogene |
| pKS1 | pIJ10770 with *lexA,* $Hyg^R$ | *lexA* was amplified using primer ks11 and ks17 followed by restriction digestion with KpnI and HindIII, and ligation into pIJ10770 cut with KpnI and HindIII | This study |
| pKS2 | pIJ10770 with *recA,* $Hyg^R$ | *recA* was amplified using primer ks18 and ks19 followed by restriction digestion with KpnI and HindIII, and ligation into pIJ10770 cut with KpnI and HindIII | This study |
| pKS3 | pIJ10770 with *lexA-3xFLAG*, $Hyg^R$ | *3xFLAG* tag was first fused to *lexA* by overlap extension PCR combining PCR obtained with primer ks11/ks12 and ks13/ks14. using primer ks11 and ks17. Final PCR product was digested and ligated into pIJ10257 cut with KpnI and HindIII | This study |
| pKS16 | pET15b(+) with *lexA,* $Carb^R$ | *lexA* was amplified using primer ks72 and ks73 followed by restriction digestion with NdeI and BamHI, and ligation into | This study |

| | | | |
|---|---|---|---|
| | | pET15b(+) cut with NdeI and BamHI | |
| pKS22 | pIJ10772 with promoter region of *recA,* $Hyg^R$ | *recA* promoter region was amplified using primer ks150 and ks151 followed by restriction digestion with HindIII and XhoI and ligation into pIJ10772 cut with HindIII and XhoI | This study |
| Pl1-B1 | Cosmid vector containing coding sequence for *recA*, $Km^R$, $Carb^R$ | | http://strepdb. streptomyces.org.uk |
| Pl2-B1 | Cosmid vector containing coding sequence for *lexA*, $Km^R$, $Carb^R$ | | http://strepdb. streptomyces.org.uk |
| pKS100 | Mutated Pl1-B1 cosmid for REDIRECT containing Δ*recA::apr,* $Km^R$, $Carb^R$, $Apr^R$ | The *recA* coding sequence (*vnz_26845*) on the cosmid vector Pl1-B2 was replaced by an oriT-containing apramycin resistance cassette, which was amplified from pIJ773 using primer ks6 and ks7. | This study |
| pKS200 | Mutated Pl2-B2 cosmid for REDIRECT containing Δ*lexA::apr,* $Km^R$, $Carb^R$, $Apr^R$ | The native *lexA* coding sequence (*vnz_27115*) was replaced by an oriT-containing apramycin resistance cassette, which was amplified from pIJ773 using the primer ks1 and ks2. | This study |

**Supplementary Table 6:** Oligonucleotides used in this study.

| Name | Sequence (5’ –> 3’) |
|---|---|
| ks1 | GCCGACGACGTGACCACCACCGCAGACAGTGCCACCATCATTCCGGGGATCCGTCGACC |
| ks2 | GAGGTGGGGTCACACCCGCCGCAGTACCGCGACGACCTTTGTAGGCTGGAGCTGCTTC |
| ks3 | TGCGTACACGCGTAAGGCAATCTGC |
| ks4 | CGCTGGCGGCCATCGACGC |
| ks5 | CGCAGCGTCGTACGTCCCGC |
| ks6 | GTGGAACCCATGGCAGGAACCGACCGCGAGAAGGCGCTCATTCCGGGGATCCGTCGACC |
| ks7 | GTCACCGGGTCAGCTCTTGACCGCGGTGGCCTTGGTCGCTGTAGGCTGGAGCTGCTTC |
| ks8 | TGTCGCCGAGTTGTCCACAGGC |
| ks9 | GGGGTGACAGCTCTTTCCCGGC |
| ks10 | ACCACCGCGATCTTCATCAACCAGC |
| ks11 | GGCGAAGCTTGGTTCCGACCCTCCTCATGGCAC |
| ks12 | TCGATGTCGTGGTCCTTGTAGTCGCCGTCGTGGTCCTTGTAGTCCACCCGCCGCAGTACCGC |
| ks13 | CGACTACAAGGACCACGACATCGACTACAAGGACGATGACGACAAGTGACCCCACCTCGCCGGC |
| ks14 | GGCGGGTACCGACCGACCGCAATCAGGTACGGAAG |
| ks17 | GCCGGGTACCTCACACCCGCCGCAGTACCG |

| | |
|---|---|
| ks18 | GCGGAAGCTTCGGCGTTGGCAGGGCATGC |
| ks19 | GCCGGGTACCCCGGTCACCGGGTCAGCTCTTG |
| ks32 | TTCGGCACGTTGTCGCTGT |
| ks72 | TCGGCATATGACCACCACCGCAGACAGTGCCA |
| ks73 | TCGGGGATCCTCACACCCGCCGCAGTACCGC |
| mb1 | TGTTCTGCGCAGCCTCAATC |
| mb2 | CTCTTCGCTGCGACGCTCTT |
| ks150 | CCGGAAGCTTACTCGCCCCCTGGTCCATGC |
| ks151 | CCGGCTCGAGCACCCGGTTTGCTTGAGTCGATCG |
| Oligonucleotides used for SPR | |
| memeFwd | GTCATCGAACGCAGGTTCGCAGT |
| memeRev | ACTGCGAACCTGCGTTCGATGACcctaccctacgtcctcctgc |
| shuffleFor | GTCAAGAAGCTGCGGTTTTCAGT |
| shuffleRev | ACTGAAAACCGCAGCTTCTTGACcctaccctacgtcctcctgc |
| recAFor | GTCATCGAACATCCATTCTCAGT |
| recARec | ACTGAGAATGGATGTTCGATGACcctaccctacgtcctcctgc |
| lexAFor_1 | GTCTTCGAAAGGTTGCGCCAAGT |
| lexARev_1 | ACTTGGCGCAACCTTTCGAAGACcctaccctacgtcctcctgc |
| lexAFor_2 | GTCCAAACACACGTTCGAGTAGT |
| lexARev_2 | ACTACTCGAACGTGTGTTTGGACcctaccctacgtcctcctgc |
| dnaEFor_1 | GTCATCGTACGTACGTTCCCAGT |
| dnaERev_1 | ACTGGGAACGTACGTACGATGACcctaccctacgtcctcctgc |
| dnaEFor_2 | GTCTTCGAACGTCCGTACGAAGT |
| dnaERev_2 | ACTTCGTACGGACGTTCGAAGACcctaccctacgtcctcctgc |
| dnaEFor_3 | GTCTTCGACCCACCCGGGTCAGT |
| dnaERec_3 | ACTGACCCGGGTGGGTCGAAGACcctaccctacgtcctcctgc |
| uvrAFor | GTCATCGAATGTGCGTGCTAAGT |
| uvrARev | ACTTAGCACGCACATTCGATGACcctaccctacgtcctcctgc |
| ruvCFor | GTCGTCCACCCCGAGAACACAGT |
| ruvCRev | ACTGTGTTCTCGGGGTGGACGACcctaccctacgtcctcctgc |
| AlkBFor | GTCCGGTTACGCTGGATCCAAGT |
| AlkBRev | ACTTGGATCCAGCGTAACCGGACcctaccctacgtcctcctgc |
